# Optimized culture of retinal ganglion cells and amacrine cells from adult mice

**DOI:** 10.1101/2020.06.16.155069

**Authors:** Yong H Park, Joshua D Snook, Iris Zhuang, Guofu Shen, Benjamin J Frankfort

## Abstract

Cell culture is widely utilized to study the cellular and molecular biology of different neuronal cell populations. Current techniques to study enriched neurons *in vitro* are primarily limited to embryonic/neonatal animals and induced pluripotent stem cells (iPSC). Although the use of these cultures is valuable, the accessibility of purified primary adult neuronal cultures would allow for improved assessment of certain neurological diseases and pathways at the cellular level. Using a modified 7-step immunopanning technique to isolate for retinal ganglion cells (RGCs) and amacrine cells (ACs) from adult mouse retinas, we have successfully developed a model of neuronal culture that maintains for at least one week. Isolations of Thy1.2^+^ cells are enriched for RGCs, with the isolation cell yield being congruent to the theoretical yield of RGCs in a mouse retina. ACs of two different populations (CD15^+^ and CD57^+^) can also be isolated. The populations of these three adult neurons in culture are healthy, with neurite outgrowths in some cases greater than 500µm in length. Optimization of culture conditions for RGCs and CD15^+^ cells revealed that neuronal survival and the likelihood of neurite outgrowth respond inversely to different culture media. Serially diluted concentrations of puromycin decreased cultured adult RGCs in a dose-dependent manner, demonstrating the potential usefulness of these adult neuronal cultures in screening assays. This novel culture system can be used to model adult neurons *in vivo*. Studies can now be expanded in conjunction with other methodologies to study the neurobiology of function, aging, and diseases.

## Introduction

*In vitro* culture of enriched neuronal populations allows for enhanced study of direct/intrinsic factors under a variety of conditions. Such cultures are commonly derived from either embryonic/neonatal tissues [1-5] or induced pluripotent stem cells (iPSCs) [6-8]. Embryonic/neonatal tissues have weak neuronal connections which allow for their isolation and culture without severe injury [9] and are particularly useful for the study of neuronal development [10-13]. iPSCs can be used to generate many neurons, which makes them an ideal choice for high throughput experiments [14]. However, recent transcriptomic data show that iPSCs are more similar to immature than mature neurons [15]. The immaturity of embryonic/neonatal and iPSC derived neurons therefore make them suboptimal for the study of age-related or inducible disorders.

As an extension of the CNS, the retina is a commonly used model to study different neuron types [16]. The retina is composed of several neuron types that are well characterized from early development through adult stages [17-27]. Within the retina, the RGCs primarily relay visual information to the brain [28]. In mice, RGCs are one of the least abundant retinal cell types, comprising only about ∼50,000 cells [29]. This paucity of RGCs makes it challenging to study cell-specific changes to RGCs in development and disease. The challenges are even greater for *in vitro* assessments, and only a few studies have successfully cultured primary adult RGCs [30-35]. In these cases, RGC neurons have typically survived only a few days *in vitro* (DIV) and displayed limited or no neurite outgrowth.

In this manuscript, we developed a 7-step immunopanning technique to isolate retinal neurons from adult mice. We successfully enriched three neuronal cell populations – RGCs and two subpopulations of ACs. After one-week of culture (7 DIV), most cells remained viable. Surprisingly, many neurons also extended neurites, many which were much longer than those previously published. Cell viability and neurite outgrowth were inversely affected by culture media conditions. Finally, RGCs treated with puromycin for 1 week showed a typical dose-response curve, suggesting that these enriched adult neuronal cultures could be used for future screens.

## Materials and Methods

### Isolation and Culture of Adult Mouse RGCs and ACs

All procedures were performed under the Association for Research in Vision and Ophthalmology (ARVO) policy for animals in vision research and approved by the Institutional Animal Care and Use Committee (IACUC) of Baylor College of Medicine. Adult C57BL/6J mice of both genders were purchased from Jackson Labs (#000664; Bar Harbor, ME). RGCs and ACs were purified from 12 week old mice utilizing a 7-step immunopanning technique modified from Barres et al. 1988 [2] and Park et al 2019 [36]. In some cases, additional RGCs were cultured from aged (>10 months) animals. Dissected retinas were enzymatically digested with 9 units/mL of papain (#LS003126; Worthington, Lakewood, NJ) for 30 mins at 34°C followed by trituration. Dissociated retinal cells were then sieved through a 20µm nylon mesh (#SCNY00020; EMDMillipore, Burlington, MA) to remove tissue clumps, resulting in a retinal cell suspension. Cell suspensions were transferred and incubated onto two subsequent negative panning plates coated with unconjugated Griffonia (Bandeiraea) Simplicifolia Lectin 1 (BSL-1) (#L-1100; Vector Laboratories, Burlingame, CA) as previously described [37], to remove monocytes and macrophages. A third negative plate coated with CD11b/c (#554859; BD Pharmingen, San Jose, CA), also removed macrophages. Cone and rod photoreceptors were eliminated with two subsequent negative selection plates coated with peanut agglutinin (PNA) (#L-1070; Vector Laboratories, Burlingame, CA) and monoclonal rat anti-mouse CD73 (clone TY/23) (#550738; BD Pharmingen, San Jose, CA), respectively. The transfer of the retinal suspension onto panning plates bound with monoclonal mouse anti-SSEA-1 (CD15)(MC480 clone) (#BD560079; BD Pharmingen, San Jose, CA) and monoclonal mouse anti-HNK-1/N-CAM (CD57)(VC1.1 clone) (#C6680-100TST; Sigma Aldrich, St. Louis, MO) allowed for the positive selections of ACs. Final panning using antibodies against Thy1.2 (CD90.2) (#MCA02R; Bio-Rad Antibodies, Hercules, CA) positively selected for RGCs. Positive panning plates of CD15^+^, CD57^+^, and Thy1.2^+^ attached cells were washed with Dulbecco’s Phosphate-Buffered Saline (DPBS) (#14287072; Life Technologies, Carlsbad, CA), 15, 20, and 25 times respectively, to eliminate non-adherent retinal cells. Adherent RGCs and ACs were treated with trypsin (1,250 units/mL) (#T9935; Sigma Aldrich, St. Louis, MO) and mechanically dissociated using a P-1000 pipette.

Brightfield images of each immunopanning step were taken on a Leica DMi8 inverted microscope (Buffalo Grove, IL) at 10x magnification to confirm the attachment of cells of interests. ACs and RGCs were stained with 0.4% Trypan Blue (#15250061; Gibco, Waltham, MA) and analyzed on the Countess II Automated Cell Counter (#AMQAX1000; Invitrogen, Carlsbad, CA) to determine the viability and yield of each cell type per retina for each isolation. Isolated RGCs and ACs were seeded onto 96-well plates (#655986; Greiner Bio-One, Monroe, SC) and 12mm coverslips (#1254582; Fisher Scientific, Hampton, NH, USA) at a density of 30,000 cells/cm^2^ coated with poly-D-lysine (#P6407; Sigma-Aldrich, St. Louis, MO, USA) and mouse laminin l (#3400-010-01; Trevigen Gaithersburg, MD). Cells were cultured in a serum-free defined media [37] containing a 1:1 mix of base media of DMEM(#11960; Invitrogen, Carlsbad, CA) and Neurobasal (#21103049; Invitrogen, Carlsbad, CA), NS21 supplement (made in house), Brain-Derived Neurotrophic Factor (BDNF) [100 ng/mL] (#450-02; Peprotech, Rocky Hill, NJ), Ciliary Neurotrophic Factor (CNTF) [20 ng/mL] (#450-13; Peprotech, Rocky Hill, NJ), and forskolin [8.4ng/mL] (#F6886; Sigma Aldrich, St. Louis, MO) (DN, 2x Tf/F, 5% CO_2_ media). Cytosine β-D-arabinofuranoside (Ara-C) [5µM] (#C6645; Sigma Aldrich, St. Louis, MO) was included in the media to inhibit the growth of glial cells [38]. Cells were cultured in a humidified 5% CO_2_ incubator at 37°C. Half of the media was exchanged for fresh media every two days.

### Immunocytochemistry

RGCs were cultured for 7 days *in vitro* (DIV) on 12mm coverslips before being fixed in 4% PFA for 15 mins at room temperature (RT) and then permeabilized in 0.1% Triton X-100 for 5 mins. Cells were then blocked for 1 hour at RT with 5% BSA and 5% normal donkey serum. Primary antibodies incubation with guinea pig anti-RBPMS (pan RGC marker) (1:250 dilution, #1832-RBPMS; PhosphoSolutions, Aurora, CO) and mouse anti-TUJ1 (neuron marker) (1:300 dilution, #MMS-435P; San Diego, CA) were performed overnight in a moist chamber at 4 °C. Cells were incubated with Cy3 conjugated secondary antibody (1:1000 dilution, #706-165-148; Jackson ImmunoResearch Laboratories, West Grove, PA) and Alexa Fluor 488 conjugated secondary antibody (1:1000 dilution, #A-21202; Thermo Fisher Scientific, Waltham, MA) for 1 hour at RT. Stained coverslips were sealed on to glass slides with Prolong Diamond antifade with DAPI (#P36971; Thermo Fisher Scientific, Waltham, MA). Six images per coverslips, in a fixed 3×2 grid, were acquired on a Leica DMi8 inverted microscope system (Buffalo Grove, IL). To determine RGC culture purity, RBPMS staining counts were compared to the overlap of co-labeled and only labeled DAPI positive stained cells. Immunocytochemistry of RGC RBPMS staining was performed from three isolations with a total cell count of n=11,687.

### Culture Condition Modifications for Adult Mouse RGC and AC cultures

Testing of growth conditions commenced after seeding isolated RGCs and ACs onto 96-well plates for one hour. Three components (base medium, trophic factors/forskolin (Tf/F) concentration, and CO_2_ concentration) of the previously described (section 2.1) serum-free defined medium culture condition were modified to determine cell viability and neurite growth. First, we varied the base media component for neuronal culture: (Neurobasal (N; see above), Neurobasal-A (NA; #10888022; Invitrogen, Carlsbad, CA), 1:1 mix of DMEM/Neurobasal (DN), 1:1 mix of DMEM/Neurobasal-A (DNA), and 4:1 mix of DMEM/water (DW)). Second, we tested the concentration of Tf/F at two different levels: “1x” concentration (BDNF [50ng/mL], CNTF [10ng/mL], and forskolin [4.2µg/mL]) and “2x” concentration (BDNF [100ng/mL], CNTF [20ng/mL], and forskolin [8.4µg/mL]). Third, we adjusted CO_2_ concentration to two levels (5% and 10%). Each permutation was tested, providing a total of 20 different culture conditions. RGCs and ACs were grown in the 20 conditions for 7 DIV with Ara-C, with 50% of the media exchanged out every other day as above. Culture conditions were analyzed according to the pooled effects of all 20 culture conditions in order to detect large effects of the tested base media, Tf/F concentrations, and CO_2_ concentrations (see *Statistical Analysis*).

### Cell Diameter, Cell Viability, and Neurite Outgrowth

Immediately after isolation and 7 DIV, RGCs and ACs were stained with the LIVE/DEAD Viability/Cytotoxicity kit (#L3224; Invitrogen, Carlsbad, CA) and nuclear stain, Hoechst 33342 (#H3570, Invitrogen, Carlsbad, CA) to determine live and dead cells and to visualize the neurite morphology of live cells. As a positive control for dead cells, cells were treated with 100% ice-cold methanol for 5 min before staining. Cells were incubated in Live-Cell Imaging Solution (#A14291DJ; Life Technologies, Carlsbad, CA) containing 2µM of calcein-acetomethoxy (calcein-AM), 1µM of ethidium homodimer-1 (EthD-1), and 20µg/mL of Hoechst 33342 for 30 mins in the dark at 37°C. Cells were then washed twice and incubated with Live Cell Imaging Solution for imaging on the ImageXpress Pico Automated Cell Imaging System (#IX Pico; Molecular Devices, San Jose, CA). For each well, images were captured at 10x magnification in a stitched 4×4 grid to create an acquisition region that covered 75% of the well. The viability, neurite outgrowth, # of neurite branches, and # of neurite processes of each cell were analyzed using CellReporterXpress’s (Version 2.5.1.0 Beta; Molecular Devices, San Jose, CA) Cell Scoring: 3 channels and Neurite Tracing built-in analyses. The diameter of a cell was calculated with the formula: 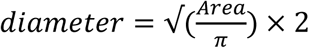, where the area of the cell was outputted by either the Cell Scoring: 3 channels [39] or Neurite Tracing analysis [40].

### Puromycin Concentration Curve

Adult RGCs were isolated and cultured on 96-wells plates in DN, 2x Tf/F, 5% CO_2_ media containing Ara-C for a 24h period before puromycin treatment. Ten-fold serial dilutions (10^4^-1ng/mL) of puromycin (#A1113803, Life Technologies, Carlsbad, CA) were applied to the RGC media for 7 DIV. Half of the media (with puromycin + Ara-C) was exchanged every other day. Viability of puromycin treated cells was determined from RGCs that were stained, imaged, and analyzed on the ImageXpress Pico Automated Cell Imaging System, as described in section 2.4 (n=3).

### Statistical Analysis

Statistical analyses were performed using Prism (GraphPad, La Jolla, CA). Two-tailed t-tests were used to compare CO_2_ concentrations and Tf/F concentrations. One-way ANOVAs using Tukey’s multiple comparisons test were used to compare among base media. Statistical significance of the experimental data was described as ^*^ = P < 0.05; ^**^ = P< 0.01; ^***^ = P< 0.001; ^****^ = P< 0.0001. Data are presented as mean ± SEM.

## Results

### RGC isolation from adult mice using a 7-step immunopanning method

We adapted our previous 4-step immunopanning method to isolate RGCs from adult mice by adding three additional steps to improve RGC purity and yield (Figure 1). To debulk the retinal suspension and allow for improved dilution and dispersion of cells in subsequent panning steps, we removed cone and rod photoreceptors with negative selection plates coated with PNA [41] and anti-CD73 [42], respectively. To further deplete ACs in order to allow for improved binding of RGCs to Thy1.2 antibody in the final step, we added a selection plate coated with anti-CD15 [43] just prior to the existing anti-CD57 plate. We did not modify the other steps (negative panning with BSL1 and anti-CD11b/c and positive selection with anti-Thy1.2). The three additional steps increased panning time by 50 minutes, resulting in a total time of 140 minutes. Despite this increased time, the new 7-step immunopanning technique produced large, circular, (Figure 2A) and healthy (Figure 2B; 91.1 ± 1.0% viable) Thy1.2^+^ cells at high yields (Figure 2B; 49,358 ± 6,655 cells per retina) consistent with our prior study [36]. Immunostaining of Thy1.2^+^ cells with RBPMS revealed that these cells are highly enriched for RGCs, reaching 85.4 ± 1.7% purity.

**Figure 1.**
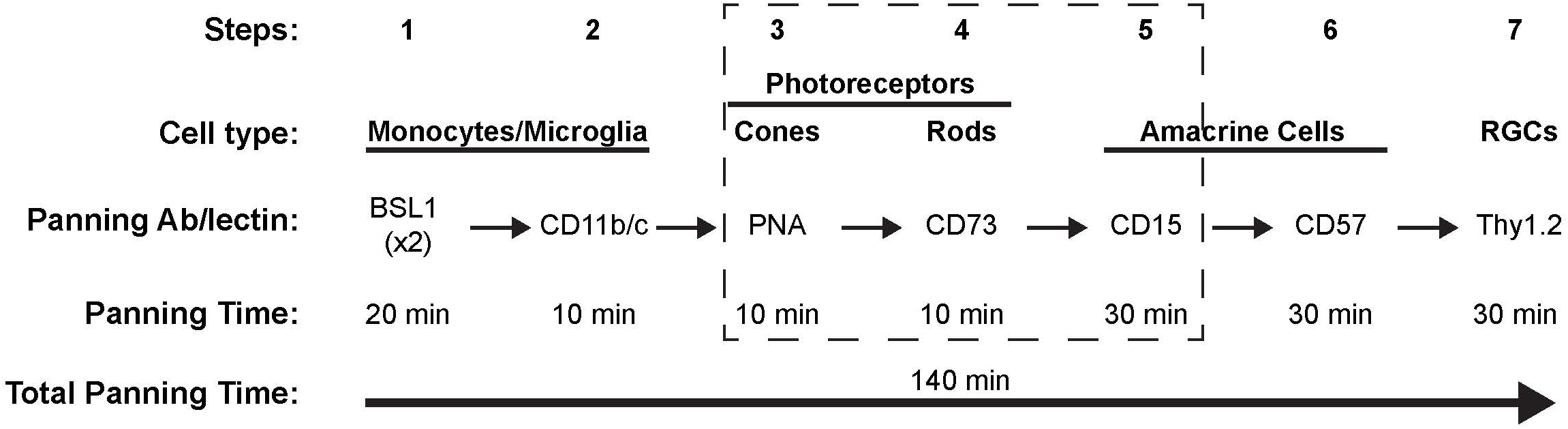
Schematic diagram of the 7-step immunopanning method to isolate adult mice ACs and RGCs. Following retinal dissociation, the cell suspension is panned through a series of plates coated with lectins and antibodies adhering cells of interest to be either negatively or positively selected. The initial four negative panning steps allow for the removal of monocytes, microglia, and photoreceptors from the retinal cell suspension. Positive panning (cells of interest to be collected) steps #5 and #6 select for ACs, while positive panning step #7 selects for RGCs. The total immunopanning time is 140 minutes. The dotted line box highlights three new steps that are improvements in the technique to isolate AC and RGCs successfully.

**Figure 2.**
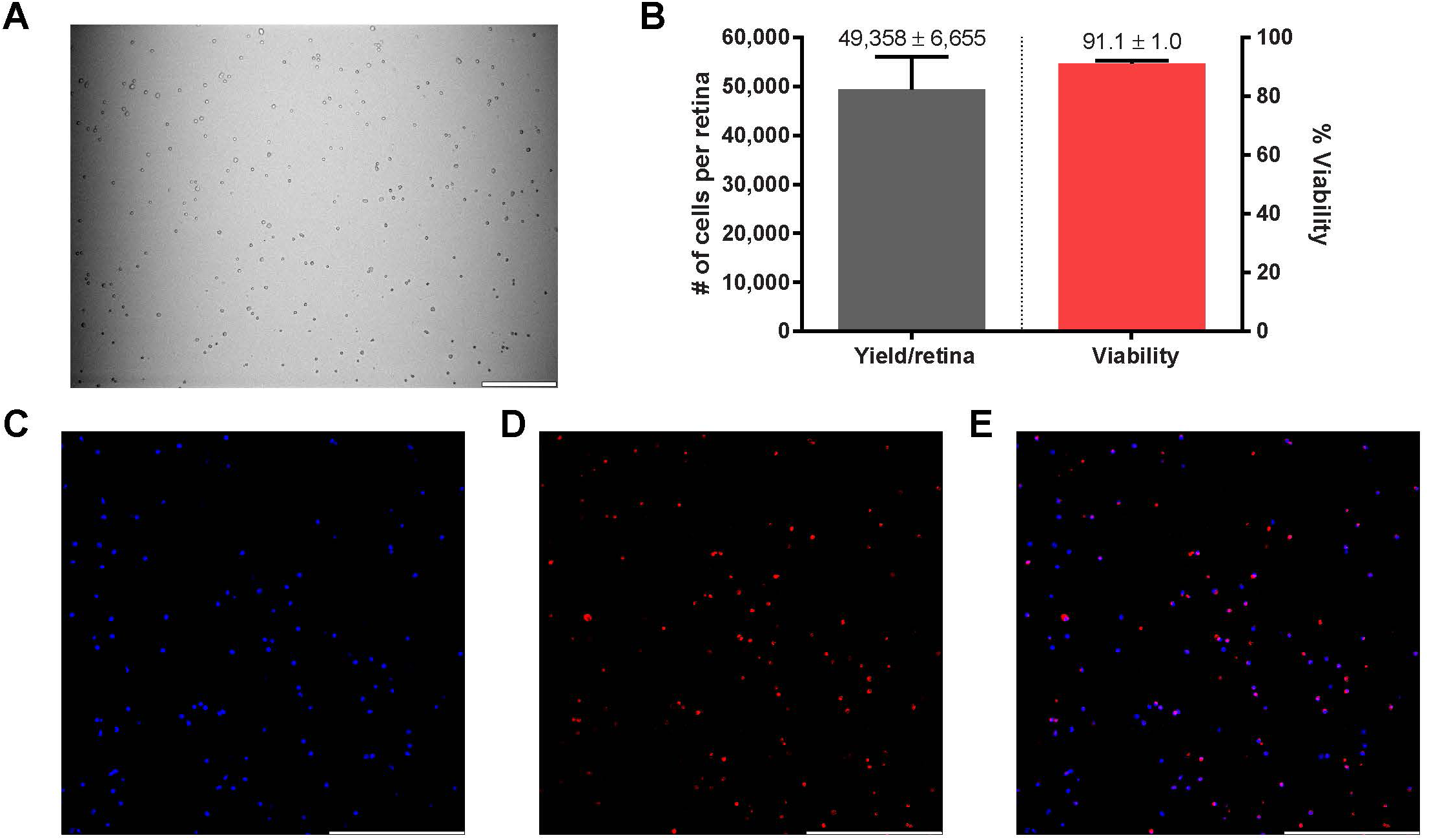
Characterization of purified RGCs isolated from adult mice. (A) Brightfield image (10x) of panning step #7 of Thy1.2^+^ adherent cells after washes. (B) The 7-step immunopanning method to isolate Thy1.2^+^ cells yields 49,358 ± 6,655 cells per retina with a 91.1 ± 1.0% cell viability (n=5). Thy1.2^+^ cells, cultured for 7 DIV, immunostained for an RGC marker, RBPMS (D), and nuclear stain, DAPI (C). RBPMS co-stained 85.4 ± 1.7% of DAPI positive cells in (E) merged image (n=3). Scale bar = 200µm.

### RGC culture and use as a possible screening assay

Enzymatic and mechanical dissociation inflicts injury and stress on cells during isolation has limited prior attempts to culture adult RGCs [44, 45]. Nevertheless, because of the high yield and viability of our isolated RGCs, we sought to determine conditions that could allow for their culture (Figure 3). RGCs were treated for 7 days according to a series of 20 possible permutations of culture conditions. These comprised 5 different base media, 2 concentrations of Tf/F, and 2 concentrations of CO_2_ (n=5; see Methods). Viability was good across all culture conditions, with an overall viability of 50.0 ± 2.3% at 7 DIV. However, Live/Dead/nuclear analysis of RGC viability revealed subtle differences (Table 1). For example, base medium DNA (34.4 ± 5.2%) significantly (p<0.05) decreased RGC viability compared to the base medium N (62.8 ± 6.6%). RGC viability was not affected by changes in Tf/F or CO_2_. Surprisingly, we found that ∼10-25% of RGCs extended neurites at 7 DIV. Neurites were either single or multi/complex (Figure 4A), in some cases reaching >500µm in length (Figure 4B). Neurites were also seen from cultured RGCs isolated from aged mice (>10 months; example in Figure 4C,D). Neurite analysis, across the same culture permutations, showed a generally improved performance of base medium DNA against all other media, in particular when compared to base medium N (p<0.001; Table 1). Thus, the base medium with the worst cell viability, DNA, had the highest percentage of live RGCs containing neurite processes (23.8 ± 1.7%). The converse was true for base medium N, which had the best cell viability but the lowest percentage of live RGCs containing neurite processes (14.1 ± 1.6%; p < 0.01). Additionally, culturing in 5% CO_2_ (20.7 ± 1.5%) resulted in a higher percentage of live RGCs containing neurite processes than 10% CO_2_ (15.0 ± 1.5%; p < 0.05). No differences were observed in the mean RGC outgrowth length, # of branches, and # of processes according to culture conditions. Thus, the choice of base medium and CO_2_ concentration appears to be a determinant of the number of RGCs that extend neurites, but not of the quality and type of these neurites.

**Table 1.** Thy1.2^+^ Culture Conditions Viability and Neurite Analyses.

|  | Viability | Neurite analysis |  |  |  |
| --- | --- | --- | --- | --- | --- |
|  | % alive | % cells w/processes | Mean outgrowth length (μm)/cell | Mean # of branches/cell | Mean # processes/cell |
| **Base Medium** |  |  |  |  |  |
| N<sup>a</sup> | 62.8 ± 6.6 | 14.1 ± 1.6 | 94.6 ± 8.3 | 1.2 ± 0.2 | 1.4 ± 0.0 |
| NA<sup>b</sup> | 54.5 ± 5.8 | 16.9 ± 1.1 | 103.4 ± 7.3 | 1.4 ± 0.2 | 1.5 ± 0.0 |
| DN | 50.5 ± 8.1 | 19.0 ± 1.8 | 108.6 ± 8.6 | 1.6 ± 0.2 | 1.6 ± 0.1 |
| DNA<sup>c</sup> | 34.4 ± 5.2 <sup>a*</sup> | 23.8 ± 1.7 <sup>a**b*</sup> | 103.1 ± 6.7 | 1.5 ± 0.2 | 1.6 ± 0.0 |
| DW | 47.9 ± 5.7 | 15.5 ± 1.4 <sup>c**</sup> | 106.4 ± 7.4 | 1.6 ± 0.2 | 1.5 ± 0.1 |
| **Trophic factors/Forskolin (Tf/F) Concentration** |  |  |  |  |  |
| 1x | 53.4 ± 4.1 | 18.2 ± 1.2 | 104.2 ± 6.1 | 1.4 ± 0.1 | 1.6 ± 0.1 |
| 2x | 51.4 ± 3.4 | 17.5 ± 1.0 | 102.3 ± 7.4 | 1.5 ± 0.2 | 1.5 ± 0.0 |
| **% CO<sub>2</sub>** |  |  |  |  |  |
| 5 | 53.7 ± 4.1 | 20.7 ± 1.5 <sup>*</sup> | 106.1 ± 9.1 | 1.5 ± 0.3 | 1.6 ± 0.1 |
| 10 | 47.4 ± 4.0 | 15.0 ± 1.5 | 100.3 ± 5.7 | 1.4 ± 0.2 | 1.5 ± 0.0 |

Data are expressed as mean ± SE.
Superscript letters are significant differences between base medium groups.
Symbols <sup>\*</sup>, <sup>\*\*</sup> are levels of significance at p<0.05 and p<0.01, respectively.

**Figure 3.**
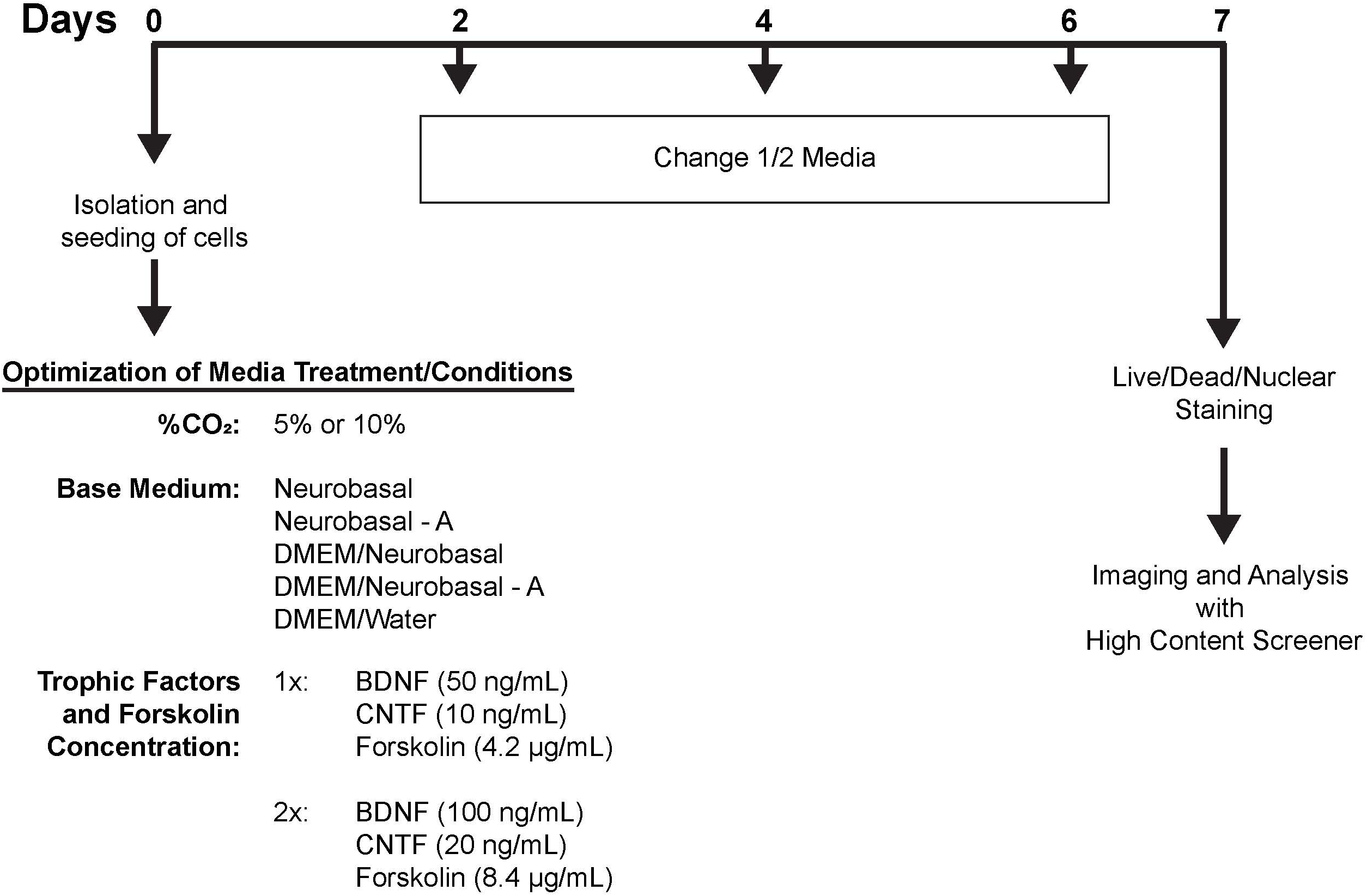
Schematic of study design optimizing culture conditions for adult mouse RGCs and ACs. Post isolation and seeding of RGCs and ACs, cells are treated with 20 different potential culture conditions for 7 days. Half the media is exchanged for fresh media containing the same components every other day. At the end of 7 DIV, cells are stained, imaged, and analyzed using a high content screener.

**Figure 4.**
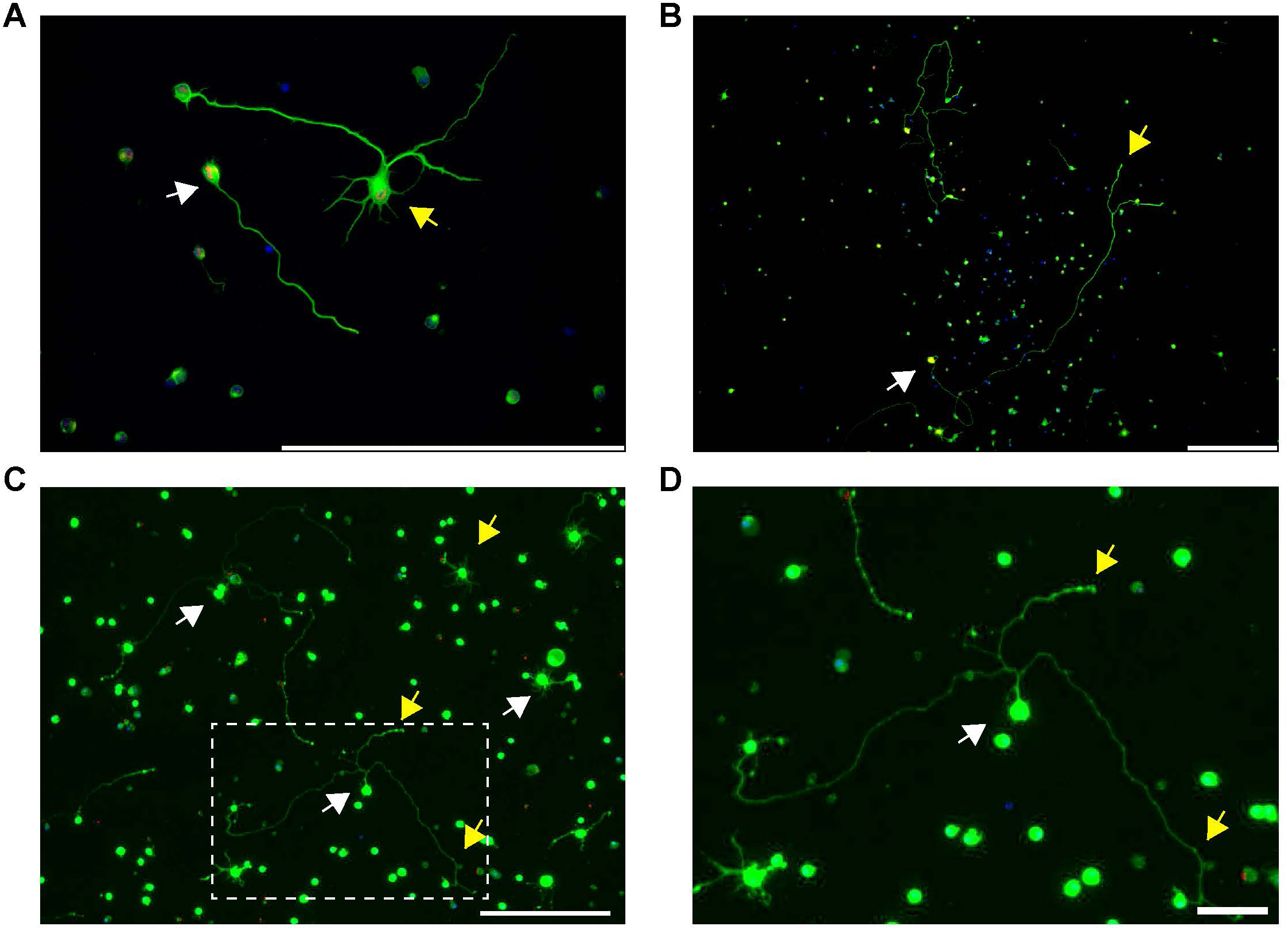
Neurite extensions from cultured adult mouse RGCs at 7 DIV. (A,B) Adult mouse RGCs immunostained for RBPMS (red), Tuj1 (green), and DAPI (blue) (Scale bar = 200µm). (A) RGCs can extend single (white arrow) and multi/complex neurites (yellow arrow). (B) Example of long (>500µm) RGC neurite extension (yellow arrow) from the soma (white arrow) (Scale bar = 200µm). (C) Live (green)/Dead (red)/nuclear (blue) staining of RGCs isolated from aged adult mice older than 10 months are viable (white arrows) and extend neurites (yellow arrows) (Scale bar = 200µm). (D) Zoom-in of the boxed area in C (Scale bar = 50µm).

Since adult RGCs in culture can be an invaluable tool for future pharmacological and genetic studies, we tested if the application of puromycin, which is widely used as a screening assay, consistently reduced the viability of cultured RGCs in a dose-dependent manner (Figure 5) [46, 47]. Indeed, the IC_50_ of the RGC puromycin kill curve was 72.9ng/mL, which is consistent with other mammalian cells used in screening assays [48], and the three trials showed minimal variability among them. Together, the 7-step immunopanning technique is gentle enough to allow for the culture of adult mice RGCs that extend neurites. Furthermore, these cultures can likely be used as an *in vitro* screening tool for studies that requires enrich populations of RGCs from adult mice.

**Figure 5.**
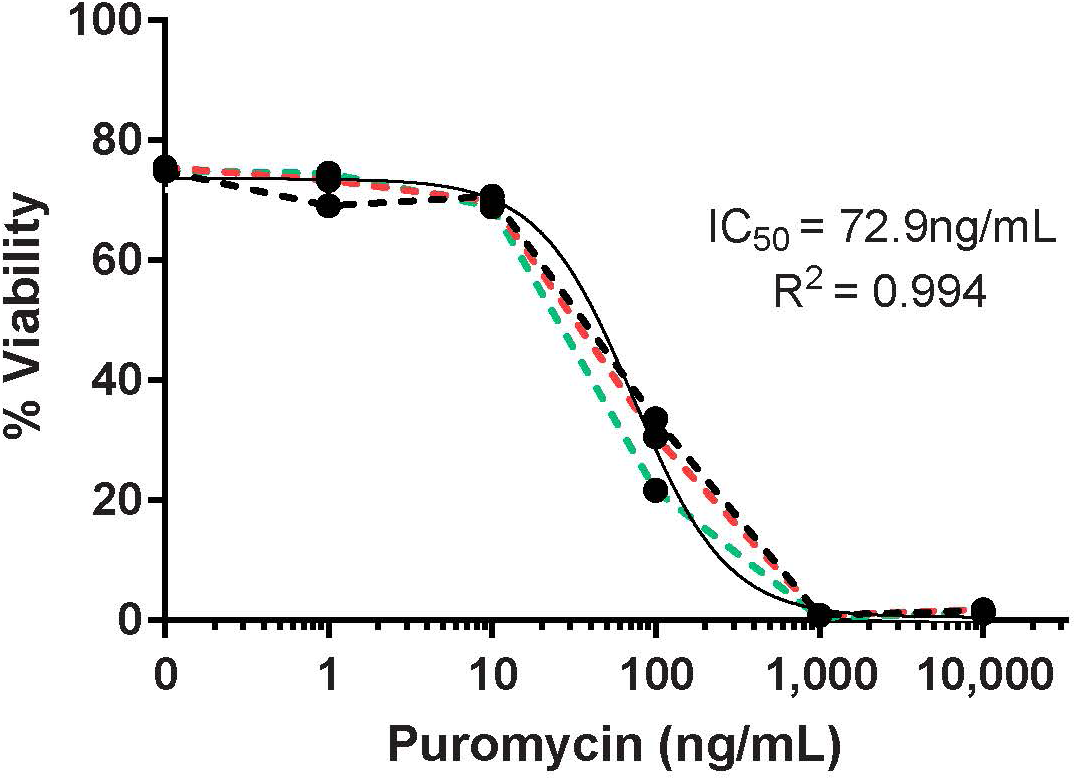
Adult RGC puromycin concentration-response curve after 7 DIV. Varying treatment concentrations of puromycin (1, 10, 100, 1,000, 10,000ng/mL) decreased RGC viability in a concentration-dependent manner. Dotted black, red, and green lines represent biological replicates (n=3). The solid black line represents the non-linear regression plot of the three replicates.

### AC culture and optimization of culture conditions

As a byproduct of the 7-step immunopanning method to isolate RGCs, two populations of ACs (CD15^+^ and CD57^+^) were isolated. Using the same approach as above, we used these populations to determine if adult ACs can also be cultured, *in vitro*. Similar to RGCs, isolated CD15^+^ and CD57^+^ cells were large and circular (Figure 6A,B), with viability greater than 75% (Figure 6C,D). The yield of CD15^+^ and CD57^+^ cells per retina were 14,717 ± 3,717 and 25,471 ± 3,447, respectively, lower than the RGC isolation yield. After 7 days, cultured adult CD15^+^ and CD57^+^ are viable and have to ability to extend neurites (Figure 6E,F). Comparable to cultured adult RGCs, base medium DNA reduced CD15^+^ viability (51.8 ± 2.1%) when compared to the best-performing base medium, which was again N (69.8±2.1%; p < 0.01; Table 2). CD15^+^ cells were significantly more viable in 5% CO_2_ (65.3 ± 2.3%), compared to cells cultured in 10% CO_2_ (57.4 ± 1.8%; p<0.01). Similar to adult RGCs, the percentage of CD15^+^ ACs with neurite processes was significantly increased in base medium DNA compared to base medium N (p<0.001). 5% CO_2_ significantly increased both the percentage of CD15^+^ cells containing neurite processes as well as the number of processes per cell (p<0.01 for both). However, CD15^+^ cells had significantly (p<0.0001) fewer branches and shorter neurites than CD57^+^ cells (compare Table 2 and 3). The two concentrations of Tf/F did not affect CD15^+^ viability or neurite extension. Finally, media conditions did not affect CD57^+^ cell viability or neurite extensions (Table 3). These results demonstrate that the 7-step immunopanning method can be adapted for successful isolation and culture of multiple cell types from a single tissue.

**Table 2.**
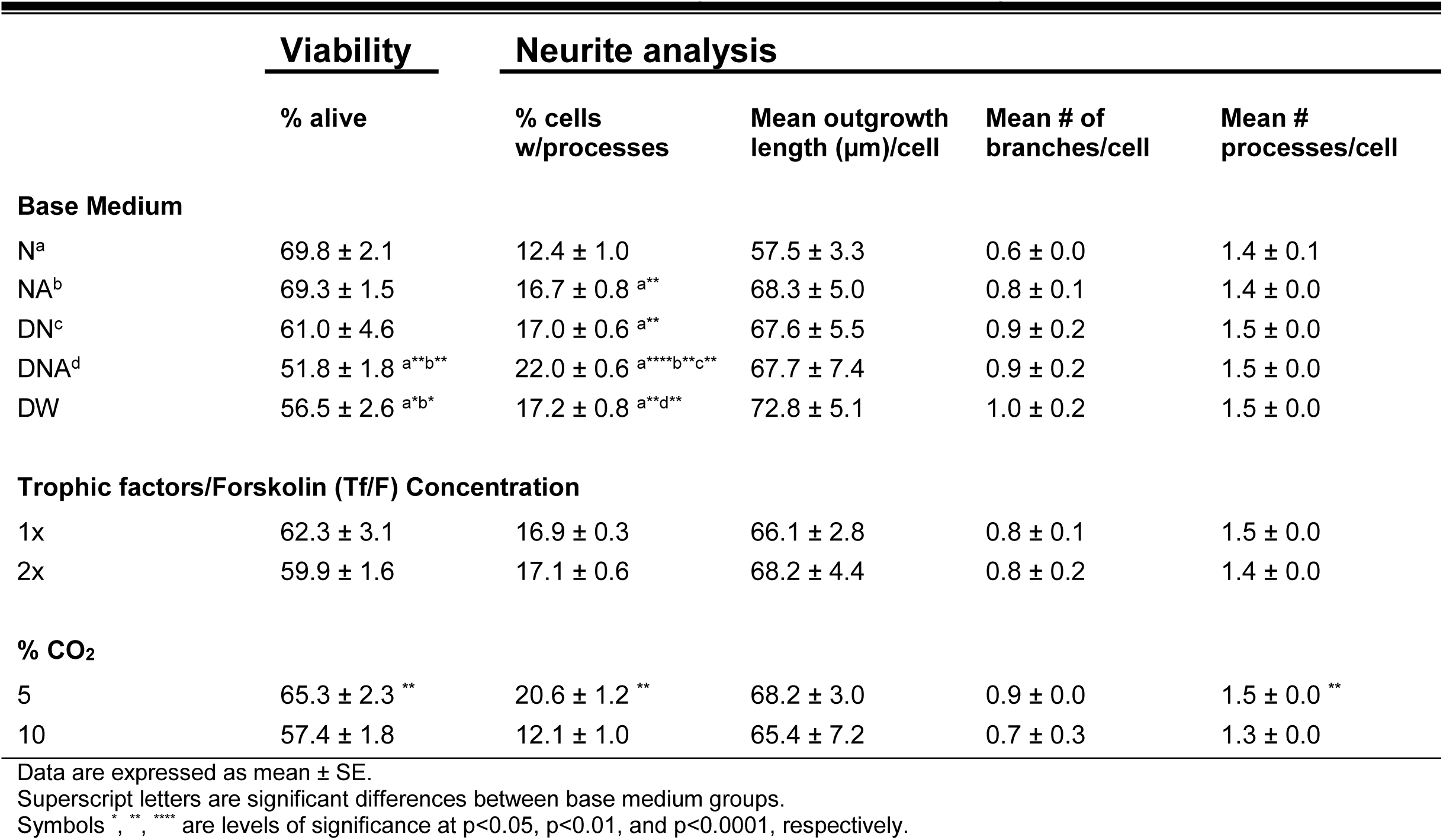
CD15^+^ Culture Conditions Viability and Neurite Analyses.

**Table 3.** CD57^+^ Culture Conditions Viability and Neurite Analyses.

|  | **Viability** | **Neurite analysis** |  |  |  |
| --- | --- | --- | --- | --- | --- |
|  | **% alive** | **% cells w/processes** | **Mean outgrowth length (μm)/cell** | **Mean # of branches/cell** | **Mean # processes/cell** |
| **Base Medium** |  |  |  |  |  |
| N | 42.8 ± 6.7 | 11.4 ± 1.0 | 95.6 ± 7.5 | 1.3 ± 0.2 | 1.5 ± 0.0 |
| NA | 39.8 ± 8.3 | 11.5 ± 0.6 | 89.0 ± 3.8 | 1.3 ± 0.1 | 1.5 ± 0.0 |
| DN | 33.8 ± 5.5 | 11.5 ± 0.3 | 77.9 ± 4.4 | 1.1 ± 0.1 | 1.5 ± 0.1 |
| DNA | 26.7 ± 5.4 | 13.6 ± 0.9 | 82.3 ± 10.7 | 1.2 ± 0.2 | 1.5 ± 0.1 |
| DW | 23.5 ± 4.8 | 11.9 ± 1.4 | 100.4 ± 15.7 | 1.6 ± 0.4 | 1.5 ± 0.1 |
| **Trophic factors/Forskolin (Tf/F) Concentration** |  |  |  |  |  |
| 1x | 38.1 ± 3.0 | 11.4 ± 0.9 | 86.5 ± 9.4 | 1.3 ± 0.2 | 1.5 ± 0.0 |
| 2x | 37.5 ± 3.3 | 12.6 ± 0.6 | 91.6 ± 5.1 | 1.3 ± 0.1 | 1.5 ± 0.1 |
| **% CO<sub>2</sub>** |  |  |  |  |  |
| 5 | 34.2 ± 2.4 | 12.3 ± 0.5 | 84.6 ± 7.9 | 1.2 ± 0.2 | 1.5 ± 0.0 |
| 10 | 40.1 ± 2.1<sup>†</sup> | 11.0 ± 1.3 | 91.6 ± 8.2 | 1.3 ± 0.2 | 1.4 ± 0.1 |

Data are expressed as mean ± SE.
<sup>†</sup>p=0.07

**Figure 6.**
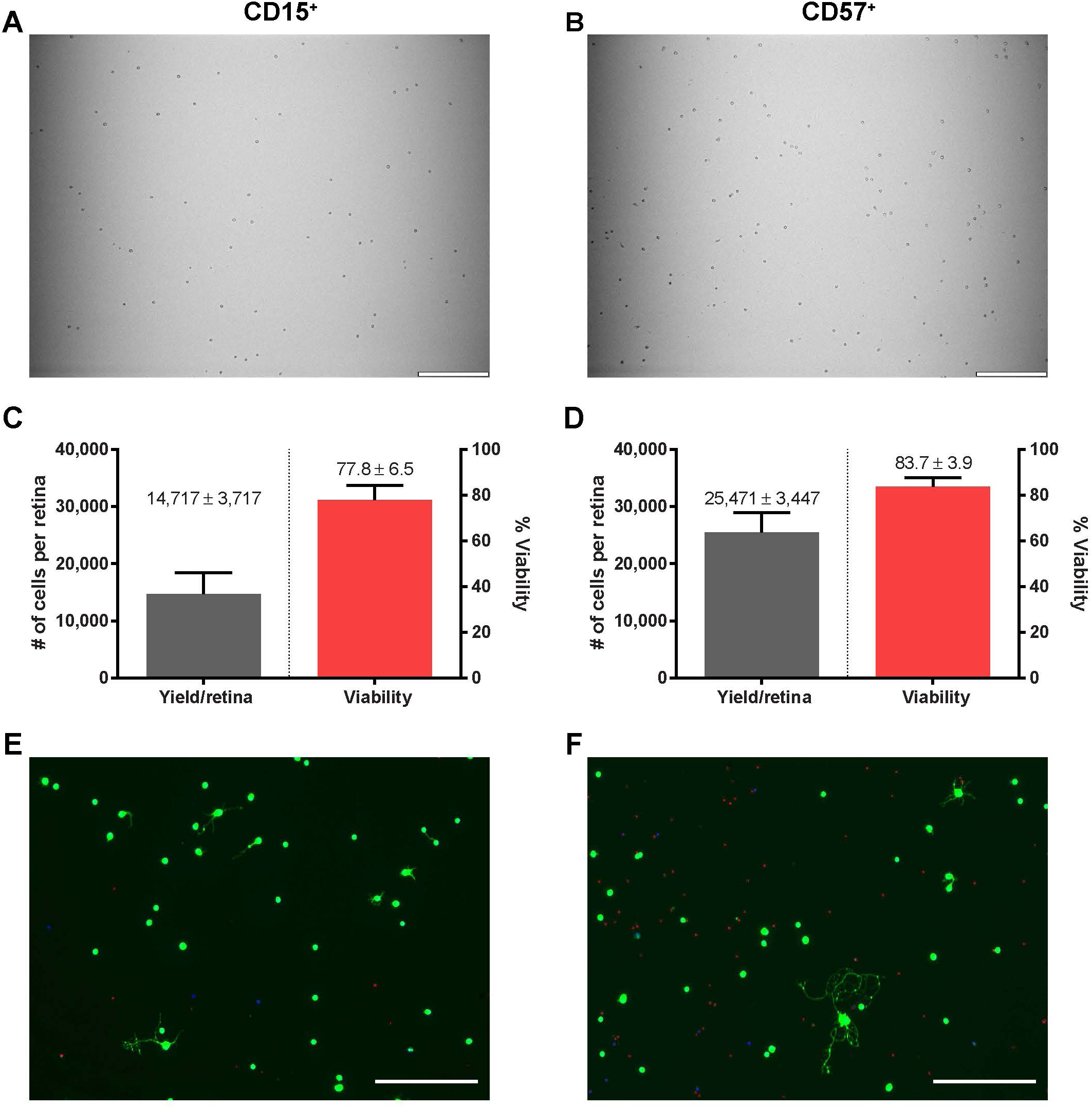
Isolation and culture of CD15^+^ and CD57^+^ ACs from adult mice. Brightfield (10x) images of attached (A) CD15^+^ and (B) CD57^+^ cells to positive panning plates after non-adherent cell washes. Collected cell yield and viability of (C) CD15^+^ and (D) CD57^+^ cells after immunopanning (n=5). Live/Dead/nuclear images of (E) CD15^+^ and (F) CD57^+^ cells after 7 DIV.

### Diameter diversity of isolated and cultured RGCs and ACs

The somas of the various subpopulations of RGCs [49, 50] and ACs [51, 52] range widely in diameter. Soma diameters of live Thy1.2^+^, CD15^+^, and CD57^+^ cells were therefore measured immediately following isolation and at 7 DIV to determine if isolations of these cell populations are diverse. The average soma diameters of recently isolated (0 DIV) Thy1.2^+^, CD15^+^, and CD57^+^ cells are 8.5, 8.7, and 8.6µm, respectively. However, all three cell populations varied widely in soma diameter, showing a range of 4-36µm with a unimodal distribution where the majority of the cell diameters range from 4-15µm (Figure 7A,B,C). When culture times for all three cell populations extended to 7 DIV, the live cell soma diameter broadened to 5-41 µm in diameter. Furthermore, instead of a unimodal distribution, all three cell populations had a bimodal distribution at 7 DIV, showing both small and large diameter somas (Figure 7A,B,C). Lastly, Thy1.2^+^, CD15^+^, and CD57^+^ cells with neurites cultured for 7 DIV have average soma diameters similar to all live cells cultured for 7 DIV. The range of the diameters of each neuron population with neurites (Figure 7D,E,F) resembles the right hump of the corresponding cell population bimodal distribution. This may indicate that the right hump contains healthier neurons compared to the left hump, which may represent dying neurons.

**Figure 7.**
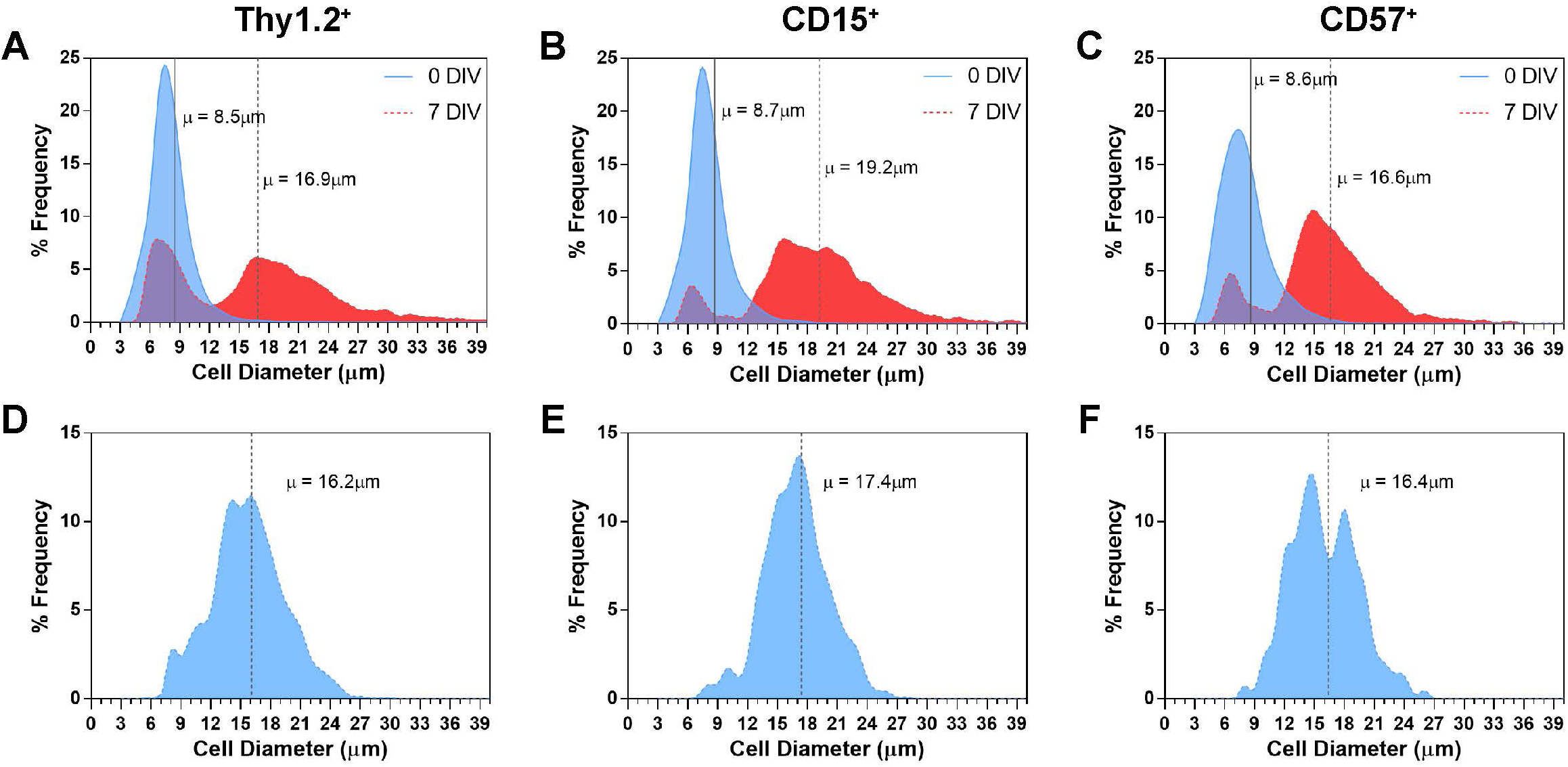
(A,B,C) Distribution frequencies of cell diameter after isolation (0 DIV; shaded blue curve) and 7 DIV (shaded red curve) for all (A), Thy1.2^+^, (B) CD15^+^, and (C) CD57^+^ cells. Vertical solid and dotted lines represent the mean cell diameters at 0 and 7 DIV, respectively. (D,E,F) Cell diameter frequencies of neurite-extending (D), Thy1.2^+^, (E) CD15^+^, and (F) CD57^+^ cells after 7 DIV, with the vertical dotted lines representing the mean cell diameter for the corresponding cell type. Thy1.2^+^: n=33,692 (0 DIV), 9,910 (7 DIV), and 2,273 (7 DIV with neurites). CD15^+^: n=21,665 (0 DIV), 5,798 (7 DIV), and 1,052 (7 DIV with neurites). CD57^+^: n=33,440 (0 DIV), 6,424 (7 DIV), and 700 (7 DIV with neurites).

## Discussion

Mature CNS neurons are post-mitotic, non-dividing cells that are extremely susceptible to injury and death, thus are limited in cell culture applications [53, 54]. Using a 7-step immunopanning technique adapted from our original protocol, we have effectively isolated RGCs from adult mouse retinas with high purity and yield. Additionally, by adjusting the procedure, we were also able to extract two subpopulations of ACs during the same isolation. The week-long cultivation of all three neuron populations revealed excellent viability. Furthermore, up to ∼25% of healthy cells extend single or multiple neurite extensions of variable lengths, with some neurites exceeding 500µm. Optimization of the culture conditions for RGCs and ACs identified inverse response differences between cell viability and neurite outgrowth. As these cells are isolated from adult tissue, this technique allows for the successful culturing of enriched populations of mature neurons, where previously purified neuronal cultures were limited to neonatal and iPSCs derived neurons.

The necessity for adult neuronal *in vitro* models in parallel with the use of other experimental models is essential to delineate further function, aging, and disease pathogenesis of CNS neurons [55]. Neurons derived from neonates and iPSCs are easily generated in large quantities and can be cultured with excellent viability with extensive neurite outgrowth [3, 6]. However, prior to isolation, these neurons are limited to genetic manipulation as well by their neuronal immaturity and epigenetic gap [56]. The successful cultivation of adult RGC and AC neurons described in this paper fills those gaps, allowing use in conjunction with a much wider range of experimental models. For example, mature neurons can be cultured from aged-matched animals correlating to the ages used in parallel with *in vivo* studies. Additionally, experimental manipulation using genetic, pharmacological, and mechanical interventions on *in vivo or ex vivo* models can be performed prior to neuronal isolation. These neurons can subsequently be cultured to be further tested for their responses to different stimuli, retaining epigenetic modifications that were gained *in vivo* before cell isolation. Thus, adult neuronal cultures are likely to be more adept at addressing questions related to the neurobiology of aging, disease, and injury.

RGCs [21, 50] and ACs [25] are both heterogeneous and include multiple subtypes. A few of these subtypes have neuroprotective and regenerative properties, which may represent the live cells in our neuronal cultures [12, 57, 58]. The broad ranges in diameters of isolated and cultured Thy1.2^+^, CD15^+^, and CD57^+^ cells, indicates diverse populations were collected. Using two well-known pan-RGC markers to first isolate (Thy1.2) [2] at the theoretical yield of mouse RGCs and validate high purity with RBPMS [19], we can assume that we have captured many subtypes of RGCs. However, a specific exogenous pan-ACs marker has not yet been identified for isolation use. In a mouse retina, CD15 is localized in two populations of ACs, separated by small and large soma diameters, resembling our bimodal cell diameter distribution (Figure 7B) [43]. Isolation of adult ACs using these two markers yields a total of 40,000 cells per retina (< 1% of the retina population), a minority of total ACs [29]. When culturing the two subpopulations of adult ACs, differences between the two cell types were exposed. Interestingly, CD15^+^ ACs and adult RGCs shared similar characteristics with regard to cell survival and neurite outgrowth according to culture conditions (see below). In contrast, mature CD57^+^ cells’ survival was non-dependent to exogenous changes in culture conditions. This may reflect the observation that CD57^+^ ACs may be resistant to neurodegeneration [12]. Further characterization of these classes of neurons and their neurites are required to determine which RGC and AC subtypes they represent in our culture.

What factors are facilitating neuronal survival and neurite outgrowth in our adult culture systems? Optimization of our culture conditions revealed inverse differences in base media on their effects on cell survival versus the percentage of cells with neurite processes in RGCs and CD15^+^ neurons. The different base media differ by osmolarity ranking in osmotic concentration as N < NA < DN = DW < DNA [59, 60]. RGCs and CD15^+^ cells exposed to lower osmolarity (N) increased in cell viability, contrary to adult hippocampal neuronal cultures. Interestingly, increasing osmolarity reduced viability but increased the percentage of cells with neurites in these two cell populations (DNA). Similar increases in neurite extensions occur with PC-12 cell (peripheral neuronal cell line) exposed to hyperosmotic conditions [61]. Vitamin B12, a component found only in N and NA base media, has been shown to protect RGCs from optic nerve transection [62]. As a superoxide scavenger, vitamin B12 may have contributed to the removal of reactive oxygen species that were propagated from the stressors of our isolation method, thus promoting the survival of cultured adult RGCs and CD15^+^ cells. Lastly, CO_2_ concentrations may affect different neuronal populations. In our cultures, both RGC and CD15^+^ cells thrived better in 5% CO_2_, while CD57^+^ cell viability improved (p=0.07) in 10% CO_2_, suggesting, CD57^+^ cells may prefer a more acidic environment. Interestingly, when comparing the two AC populations, unlike for CD15^+^ cells, culture conditions had little effect on CD57^+^ cell viability and neurite outgrowth. Taken together, our results suggest that the conditions for cell survival versus neurite outgrowth can differ in the same neuronal population, and therefore having a single target therapy for a neurodegenerative disease or injury may not be beneficial for both neuronal survival and neurite regeneration. This is particularly important when considering the treatment of human diseases for which injury is ongoing, such as glaucoma. Similarly, a single culture condition may not be optimal for the use of all neuronal types. Therefore, the optimization of culture conditions based on the experimental question is important when studying and screening for neuronal survival, regeneration, phenotype, and physiology.

Cultures of enriched neuronal populations from adults can help us better understand the complexity of the CNS. Adult neuronal cultures can be used as screening tools (Figure 5) and co-cultures can be performed to study specific interactions between mature neurons and with other non-neuronal cell types. Additionally, these cultures can be expanded with other experimental methods as a tool to delineate biological relevant questions of neurons that were previously possible only in mitotic somatic cells. The increased versatility and capabilities of adult neuronal cultures complement the various strengths of neuronal cultures using neonatal cells and iPSCs, and will be particularly helpful to further delineate neuronal function in aging and disease in future studies.

## Acknowledgments and Funding

This project was funded by research awards from the Clayton Foundation for Research (Houston, TX) and the National Institutes of Health (R01 EY-025601) to BJF, and a department award to Baylor College of Medicine from Research to Prevent Blindness (New York, NY).

## Competing Interests

None.

## Notes

### Competing Interest Statement

The authors have declared no competing interest.

